# incR: a new R package to analyse incubation behaviour

**DOI:** 10.1101/232520

**Authors:** Pablo Capilla-Lasheras

## Abstract

Incubation represents a life stage of crucial importance for the optimal development of avian embryos. For most birds, incubation poses a trade-off between investing in self-maintenance and offspring care. Furthermore, incubation is affected by environmental temperatures and, therefore, will be likely impacted by climate change. Despite its relevance and readily available temperature logging methods, avian incubation research is hindered by recognised limitations in available software. In this paper, a new quantitative approach to analyse incubation behaviour is presented. This new approach is embedded in a free R package, `incR`. The flexibility of the R environment eases the analysis, validation and visualisation of incubation temperature data. The core algorithm in `incR` is validated here and it is shown that the method extracts accurate metrics of incubation behaviour (e.g. number and duration of incubation bouts). This paper also presents a suggested workflow along with detailed R code to aid the practical implementation of `incR`.

## Introduction

Incubation represents a crucial life stage for egg-laying vertebrates, of which birds are a paramount example. Fine control of incubation is essential and has deep ecological and evolutionary implications, notably for developing offspring but also for their parents (Conway and Martin 2000, Durant et al. 2013). For embryos, the thermal environment that the incubating individual provides is essential for successful development. Suboptimal incubation temperatures can lead to delayed embryonic growth (Hepp et al. 2006, Nord and Nilsson 2011), hormonal and immune changes (Ardia et al. 2010, DuRant et al. 2014), and long-term survival consequences (Berntsen and Bech 2016). However, incubating individuals need to divide their time budget between incubation and self-maintenance (e.g. foraging) and, therefore, they allocate time to each activity according to prevalent ecological conditions (e.g. ambient temperatures (Coe et al. 2015) or food availability (Londoño et al. 2008)). Despite a long standing scientific interest in incubation, we are still elucidating subtle ecological causes and consequences of variation in this behaviour (Durant et al. 2013, Smith et al. 2015, Bulla et al. 2016) which may have important practical implications, for example, in a context of global climate change (Griffith et al. 2016).

The study of avian incubation is nowadays fuelled by recent technological advances (see Smith et al. 2015). In particular, the use of iButtons® (Maxim Integrated) and probed Tinytags (Gemini Data Loggers) allows researchers to measure incubation temperature as frequently as every second for long periods of time with minimal disturbance. These technologies have the potential to expand the range of species and scientific questions that researchers can address. However, the amount of data collected is usually much larger than it was traditionally available and several analytical hurdles must be overcome.

Before answering biological questions about incubation patterns, the observer needs to summarise the data and effectively reduce them to a few variables that can be correlated with a set of predictors of interest. For example, number of incubation bouts and their duration are popular metrics in avian studies (Cooper and Voss 2013). The first software for the analysis of incubation temperatures was released more than 10 years ago: Rhythm (Cooper and Mills 2005). The benefits of Rhythm were immediate as it allowed the automated differentiation between time periods when eggs were being incubated (Cooper and Voss 2013, Coe et al. 2015). This software made fast and objective an otherwise time-consuming and subjective activity. However, in a time when incubation data collection is easier than ever before, Rhythm lacks much of the flexibility required for the handling of big data sets. Rhythm also has limited analytical and graphical capabilities, which are a desire when thousands of temperature records may be available. However, apart from Rhythm, no other specialised software is currently available to analyse incubation temperature data.

To overcome these difficulties, I have developed a new R package, `incR`. This package provides a suite of R functions that i) prepare and format a raw temperature time-series (via the `incRprep` and `incRenv` functions), ii) apply an automated algorithm to score incubation (`incRscan`), iii) plot the data (`incRplot`) and iv) calculate biologically relevant metrics of incubation (e.g. number of incubation bouts) (Figure 1). Users can apply the whole pipeline or use any of the components of `incR` separately. `incR` takes advantage of the flexibility in data handling and graphical capabilities offered by R. I first explain the workflow of `incR` and its automated algorithm to score incubation. Then, I use video-recordings of incubating blue (*Cyanistes caeruleus*) and great tit (*Parus major*) females along with incubation temperature data to validate the automated algorithm. I further show how `incR` can accurately calculate several metrics of incubation behaviour. Finally, I discuss the general application of this new method and its potential limitations. A stable version of the package is available on CRAN (v 1.1.0) and a development version can be found on GitHub (v 1.1.0.9000. https://github.com/PabloCapilla/incR).

**Figure 1.**
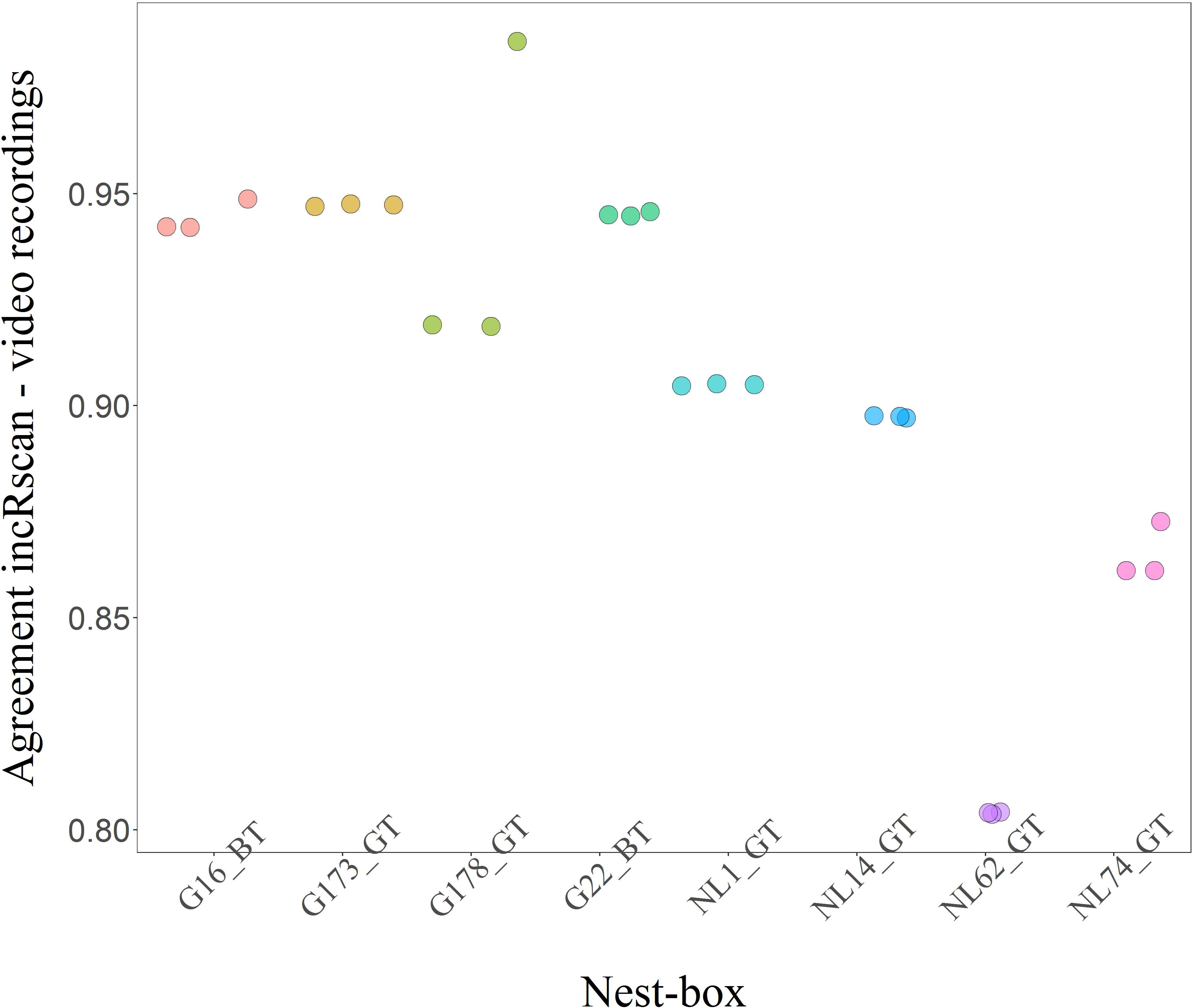
`incR` workflow and visualisation of corresponding analysis of nest temperature data at each step of the workflow. After the user collates information from a single nest, `incR` can be used. `incRprep` prepares raw data time series for the pipeline (1) and `incRenv` adds environmental temperatures to the initial data table (shown as green lines in the plot 2). `incRscan` classifies data points into absence (purple) or presence (light red) of the incubating individual in the nest (3). From a sequence of 0’s and 1’s calculated by `incRscan`, `incRbouts`, `incRatt`, `incRact` and `incRt` extract information about on/off-bouts, nest attendance, start and end of activity and averaged nest temperatures for customised time windows. `incRplot` can be used to visualise the results of `incRscan` and produce the graph shown in panel 3.

### incR workflow

The method implemented in `incR` exploits variation in nest (incubation) and ambient temperature to calculate the presence or absence of an incubating individual in the nest. Ambient temperature data are ideally collected near the nesting site but can also be obtained from web-based sources if the latter is not available. Code and advice to replicate the analyses presented here can be found in Appendix 1 and 2, the package documentation (https://cran.r-project.org/web/packages/incR/incR.pdf) and in a package vignette (accessible in R via: `browseVignettes(“incR”)`). Additionally, `incR` is distributed with an example data set that can be explored to understand data structure and the use of each `incR` function. For details to install the package, visit: https://github.com/PabloCapilla/incR

#### Data preparation: incRprep and incRenv

To start working with `incR`, the user needs to have a file with temperature and time information for a single nest under study. This file should consist of at least two columns: date-time and temperature values. Once this initial file is prepared, the first step in the pipeline is performed by `incRprep`, which simply prepares the dataset for other pipeline components. Then, `incRenv` can be used to automatically assign environmental temperature to every incubation temperature observation, information required by `incRscan` to score incubation (Figure 1). `incR` is distributed with sample data and, therefore, the user can check the data structure needed to start the pipeline.

#### Automated incubation scoring: incRscan

The algorithm implemented by `incRscan` exploits changes in nest temperature that arise from the behaviour of the incubating adult considering the difference between incubation (i.e. temperature in the nest cup) and environmental temperatures (see Table 1 for definitions of terms used throughout the paper).

**Table 1.** Glossary of terms used in this manuscript.

| Term | Type | Description | Chosen by the user? |
| --- | --- | --- | --- |
| Incubation temperature or nest temperature | - | Temperature inside the nest cup at any given time point. By this term I refer to a variable value that depends on whether, and for how long, the incubating individual is in or out the nest | - |
| Environmental temperature | - | Air temperature outside nest | - |
| incR_scan | R function | Calculates presence or absence of the incubating individual in the nest based on nest and ambient temperature variation | - |
| temp.diff.threshold | incR_scan argument | Difference allowed between nest and environmental temperatures | Yes |
| lower.time | incR_scan argument | Start of a time window when the incubating individual is assumed to be in the nest | Yes |
| upper.time | incR_scan argument | End of a time window when the incubating individual is assumed to be in the nest | Yes |
| sensitivity | incR_scan argument | Reduction in off-bout threshold when nest temperature is close to environmental temperature | Yes |
| maxNightVariation | incR_scan argument | Maximum variation allowed in the lower.time – upper.time window. It controls for big drops in temperature within this temporal window (i.e. night-time incubation off-bouts) | Yes |
| maxDrop | Internal calculation in incR_scan | Maximum drop in temperature between two consecutive time points within the lower.time – upper.time window | No. Calculated and reported by incRscan |
| maxIncrease | Internal calculation in incR_scan | Maximum increase in temperature between two consecutive time points within the lower.time – upper.time window | No. Calculated and reported by incRscan |

Four possible situations broadly exist regarding the change in nest temperature after the incubating individual enters (on-bout) and leaves (off-bout) the nest. These four scenarios are classified as follows: 1) incubation off-bout when nest temperature is high (close to maximum incubation temperature); 2) incubation on-bout when nest temperature is high (close to maximum incubation temperature); 3) incubation off-bout when nest temperature is low (close to environmental temperature); 4) incubation on-bout when nest temperature is low (close to environmental temperature). See Figure S1 for a visual representation of these four scenarios. Cases 3 and 4 are especially sensitive to the assumption that environmental temperature is lower than maximum incubation temperature (see Results and Discussion). The change in nest temperature that is expected after an incubation on-/ off-bout differs across the four scenarios.

Assuming that environmental temperature is normally lower than maximum incubation temperature, in scenario 1, when the incubating individual leaves the nest, a sharp drop in nest temperature is expected to follow (Off-bout(1) in Figure S1). At this point, any increase in nest temperature would mean that the bird has returned to the nest (scenario 2. On-bout(2) in Figure S1). If an off-bout occurs when nest temperature is close to the environmental temperature (scenario 3), the decrease in nest temperature after the event would be small (Off-bout(3) in Figure S1). When a long off-bout brings nest temperature close to the environmental one, an incubation on-bout would be reflected in a large increase in nest temperature (scenario 4. On-bout(4) in Figure S1).

These four scenarios represent simplified extremes in a spectrum of possible situations but they illustrate the general principle. To explain the analytical approach in more practical terms, I here describe the analysis of one day of incubation (day 1), using the terminology employed in the R package (Table 1).

For every time point in the incubation time series, `incRscan` calculates the difference between nest and environmental temperatures. Then, these differences are compared against the value of `temp.diff.threshold` (Table 1), determining whether scenarios 1 and 2 or 3 and 4 (see above) are applicable for a given time point. Two cases are possible: i) nest temperature is higher than environmental temperature by more than `temp.diff.threshold` degrees; or, ii) nest temperature is within `temp.diff.threshold` degrees of the environmental one.

Comparing the change in nest temperature between consecutive temperature recordings against temperature thresholds, `incRscan` determines whether the incubating individual is in the nest or an off-bout has occurred. Rather than having a fixed threshold for the entire analysis, a flexible threshold value is applied among days. Within days, the threshold to detect off-bouts can also change controlled by `temp.diff.threshold` and `sensitivity` (i.e. to accommodate changes in cooling rates between scenarios 1/2 and 3/4 – see below). No threshold choice is required from the user but they are calculated by `incRscan` for each day of analysis. To accomplish this, the user needs to specify some period of the 24-hour cycle when an incubating bird can be assumed to be incubating eggs in its nest. This time window is controlled by the arguments `lower.time` and `upper.time`, representing the start and end of the time of day (for instance, for diurnal bird species this period can be set at night, when the incubating individual rests in the nest). Within this time window, the maximum decrease in nest temperature between pairs of consecutive points is calculated and set as a threshold for incubation off-bouts (hereafter, `maxDrop`) for scenario 1. Assuming that nest temperatures are above environmental values, `maxDrop` is thought to effectively represent the maximum drop in temperature associated with periods when the incubating individual is in the nest. The threshold for incubation off-bout in situation 3 must be lower than in scenario 1 (i.e. when nest temperature is close to environmental temperature); thus, the argument `sensitivity`, that must be specified by the user (taking values from 0 to 1), allows for such reduction, setting the off-bout threshold in scenario 3 as `maxDrop` X `sensitivity`. Similarly, `maxIncrease` is defined as the maximum increase in temperature between pairs of consecutive points within the `lower.time` -`upper.time` window and is set as a threshold for incubation on-bouts in scenario 4. Any increase in nest temperature in scenario 2 would mean an incubation on-bout. Note that `maxDrop` and `maxIncrease` do not need to be chosen by the user but are calculated by `incRscan` for every day of analysis and reported in an R object named `incRscan_threshold`. See Appendix 1 and 2 for a practical example.

Once these thresholds are set, temperature differences between successive pairs of data points throughout the day and between `upper.time` and `lower.time` are calculated. These values are sequentially compared with the value of `maxDrop` and `maxIncrease`, following a set of conditions:

For scenario 1 and 2,

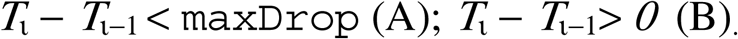

For scenario 3 and 4,

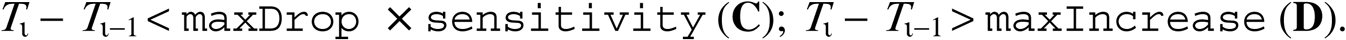

*T*_*ι*_ − *T*_*ι*−*1*_ being the *i*^th^ and *i-1*^th^ temperature recordings from *i*=2 to *i*=*I* (*I* being equal to the total number of daily data points evaluated). Off-bout periods are, then, defined between T_*i’*s_ satisfying A or C and the closest subsequent situation in which T_*j*_, when *i*<*j*, satisfies B or D. On-bout periods start after an off-bout finishes and last until A or C is fulfilled again.

This algorithm can be sensitive to highly variable temperatures or marked drops in temperature within the `lower.time` -`upper.time` window. To make `incRscan` conservative and robust against these two potential sources of error, whenever |*`maxDrop`*|`maxNightVariation` is fulfilled for a particular day of study, the value of `maxDrop` and `maxIncrease` of the previous day of incubation is instead used. `maxNightVariation` represents the maximum drop in temperature allowed in a period of constant incubation (i.e. within the `lower.time` -`upper.time` window). When this value is set too high, real off-bouts will be missed by `incRscan`.

The result of this algorithm is a temporal sequence of 0’s and 1’s representing on-bouts (1’s) and off-bouts (0’s). Using these sequences, other functions within `incR` can be used to infer incubation behaviour.

#### Additional functions to visualise nest temperatures and extract biological metrics of incubation

Regardless of whether or not `incRscan` has been used to score incubation, the `incR` package offers a suite of functions that can be applied to any binary time-series representing incubation. The current package version (1.1.0) allows the user to visualise the results of `incRscan` (`incRplot` generates a plot similar to graph 3 in Figure 1 and Figure S1), calculate onset and offset of daily activity (`incRact`), percentage of daily time spent in the nest (`incRatt`), number and average duration of on/off-bouts per day as well as individual off-bout duration and timing (`incRbouts`) and nest temperature mean and variance for a customised time window (`incRt`). The implementation of these functions is straightforward as they only require a variable with binary data for on and off-bouts. These data are provided by `incRscan` under the column name `incR_score`. The function argument `incubation.vector` in `incRact`, `incRatt`, `incRbout` and `incRt` allows the user to manually specify the name of the column with binary data for incubation scores (see Appendix 2 and package documentation in R).

### Validation of incR using temperature and video-recording data

To show the potential of this approach to yield meaningful metrics of incubation, I assessed the performance of the core functions in `incR`. First, I carried out a sensitivity analysis in which I evaluated the accuracy of `incRscan` over different values of its main arguments. I then chose the optimal values for these arguments and showed that the combination of `incRscan` and other `incR` functions can yield accurate measures of incubation behaviour. I applied the whole pipeline to incubation temperatures collected using iButton® devices. For the same incubation events, I used video footage of these nests to visually score incubation and then compared these results to the automatic algorithm implemented in `incRscan`.

#### Field protocol and incubation data collection

Incubation temperatures were recorded during 2015 and 2016 using iButton® devices in two blue tit and six great tit clutches. Blue tit data came from an urban and suburban population in Glasgow city (n = 2 clutches; 55° 52.18’N 4° 17.22’W and 55° 9’N 4° 31’W) (Pollock et al. 2017), whereas great tit incubation data were recorded in an oak forest at the Scottish Centre for Ecology and the Natural Environment (n = 2 clutches; SCENE, 56° 7.73’N 4° 36.79’W) (Pollock et al. 2017) and in a mixed forest (dominated by oak, birch and pine trees) near the Netherlands Institute for Ecology (NIOO) (n = 4; ∼52° 7’ N 6° 59’ E) (Spoelstra et al. 2015). Each iButtons® was wrapped in a piece of black cloth and placed in the nest cup, above the lining materials and among the eggs. Nest temperatures were recorded by iButtons^®^ every 2 or 3 minutes. Video cameras inside the nest-boxes were used to monitor individual females and visually score incubation (see Pollock et al. 2017 for a general explanation about video camera deployment). In total, 12 days of incubation were completely or partially monitored using both iButtons^®^ and recording cameras. Environmental temperatures for the same period in Scotland were recorded using iButtons^®^ placed outside nest-boxes. For the Dutch clutches, environmental temperatures from a weather station approximately 18 Km away from the nest-box population were used. Data from the iButtons^®^ were downloaded in the field using portable devices and a single file per nest was compiled in preparation to use `incR`.

#### Data analysis

Using video footage, I determined whether or not the incubating female had been present in the nest at every iButton^®^ temperature time point. After preparing incubation temperature data using `incRprep` and `incRenv`, I applied `incRscan` to score incubation and compared its results to the footage-based scoring. I tested the performance of `incRscan` to changing values of its three key arguments, i) `maxNightVariation` (testing values from 0.5 to 10 every 0.5), ii) `sensitivity` (from 0 to 1 every 0.1) and iii) `temp.diff.threshold` (from 0.5 to 10 every 0.5) (see Table 1 for definitions). When testing one argument, the others were kept to default values of 1.5, 0.15 and 3 for `maxNightVariation`, `sensitivity` and `temp.diff` respectively. This approach assumes that there are no interacting effects between parameter values. However, as a preliminary step in the analysis, I confirmed that that was the case. Therefore, I present here a 1-dimensional grid search (i.e. varying values of one parameter while keeping the others fixed to a given value).

`lower.time` and `upper.time` were always fixed to 10 p.m. and 3 a.m (night time). For every test, I calculated the percentage of correctly scored incubation time points. After selecting the best-performing combination of argument values (i.e. highest percentage of agreement between `incRscan` and video footage), I compared daily incubation attendance (i.e. percentage of time spent in the nest), number of daily off-bouts and mean daily off-bout duration between `incRscan`-based and video footage-based incubation scores. I present Pearson’s correlations coefficients between the two metrics. `incR` functions, statistical tests and graphical illustrations (apart from the left-hand side of Figure 1) were produced in R version 3.4.4 (R Core Team 2018). Detailed practical guidelines to use `incR` and reproduce the validation shown in this manuscript can be found in Appendix 1 and 2 as well as in the package’s vignette (accessible in R via: `browseVignettes(“incR”)`).

## Results and Discussion

Within nest-boxes, changing values of `maxNightVariation` did not affect the performance of `incRscan`. Similar results were found for `sensitivity` and `temp.diff.threshold`, with only analysis of data from one nest-box being markedly affected by changes in these arguments (Figure S2A-C). It is important to note that when `maxNightVariation` is set to a very low value (effectively not allowing for much temperature variation in the `lower.time` -`upper.time` time window) `incRscan` fails to yield any result as no temperature threshold would be available. This result can be seen in Figure S1A: when evaluating `maxNightVariation` equal to 0.5°C, data from only two out of eight nest-boxes were extracted by `incRscan.`.

Consistent variation in `incRscan` best-performing argument values was found among nest-boxes (Figure 2), suggesting that differences in, for example, iButtons^®^ deployment may be affecting the accuracy of the `incRscan` algorithm. This potential effect has been qualitatively suggested before (Smith et al. 2015) and highlights the importance of collecting high quality data in the field. However, the percentage of agreement was always high (> 80%, Figure 2). Highly consistent results were found within nest-boxes with marked among-box variation with only one exception (nest-box G178_GT, Figure 2) in which setting `maxNightVariation` to 4°C improved the percentage of agreement compared to that found with the default value (3 °C). The general pattern across the eight nest-boxes is that values above 1.5°C for `maxNightVariation` give the highest accuracy (90.27%. Figure S1D). Similarly, values below 0.3 for `sensitivity` (90.27%) and a `temp.diff.threshold` value of 4°C (91.16%) were found to be the most accurate argument choices (Figure S2E-F).

**Figure 2.**
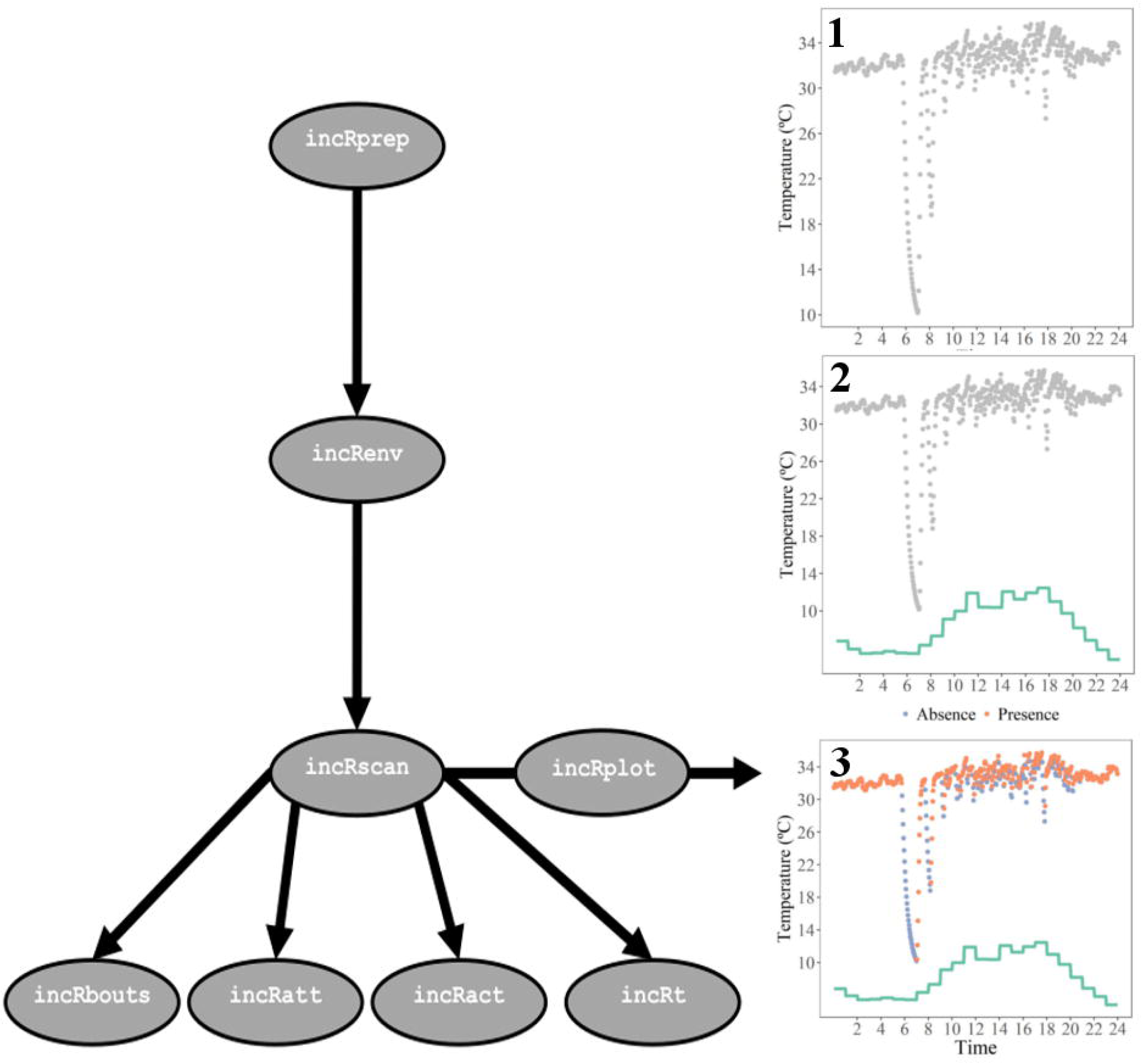
Percentage of agreement between `incRscan` and video-footage across eight different nest-boxes. Colour codes represent individual nest-boxes and each point within nest-box illustrates the percentage of agreement for each of the three 1-dimensional grid searches, after the best values were selected for `maxNightVariation`, `sensitivity` and `temp.diff.threshold`. Consistent results are found within nest-boxes with one exception (G178_GT) in which setting `maxNightVariation` to 4°C improved the percentage of agreement compared to that of the default value. Points are slightly offset in the x axis to aid visualisation of overlaying points.

Given these results, I set the parameters to their overall optimal values of 1.5°C, 0.25 and 4°C for `maxNightVariation`, `sensitivity` and `temp.diff.threshold` respectively, yielding a percentage of agreement across nest-boxes of 91.16% (maximum = 98.56%; minimum = 80.42). With these argument values, attendance calculated based on video footage and inferred by `incRscan` showed a Pearson’s correlation coefficient of 0.992 (t = 24.81, p < 0.0001, 95% confidence interval = 0.971-0.998. Figure 3A). Likewise, the algorithm in `incRscan` was able to provide accurate off-bout information (Figure 3B & 3C). `incR`-estimated off-bout number and mean daily off-bout duration were highly correlated with real off-bout number and duration as extracted from video footage (for off-bout number: r = 0.972, t_10_ = 13.04, p < 0.0001, 95% confidence interval = 0.900-0.992; for daily mean off-bout duration: r = 0.996, t_10_ = 34.69, p < 0.0001, 95% confidence interval = 0.985-0.999).

**Figure 3.**
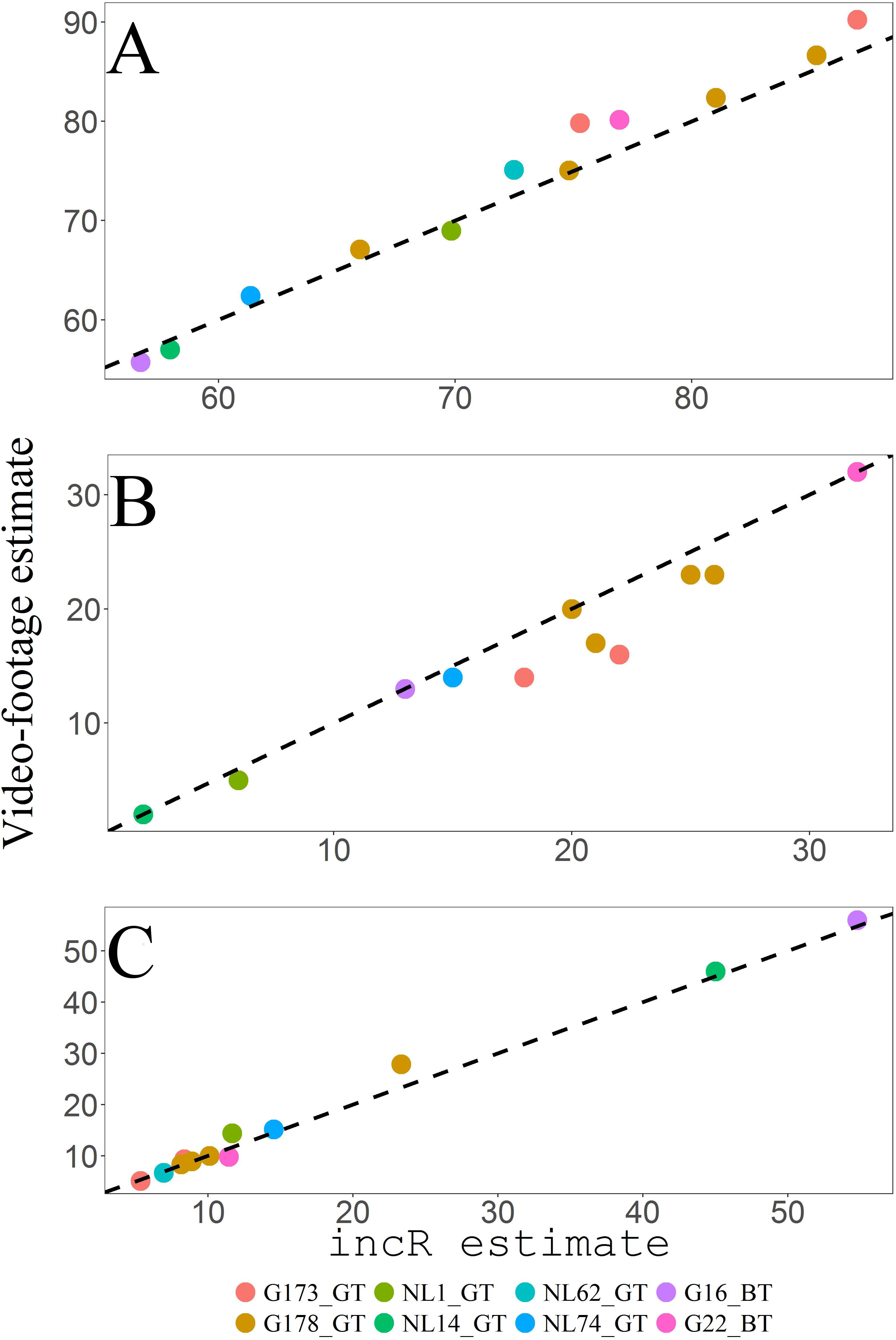
Correlations between video-footage and `incR` estimates of incubation attendance (percentage of daily time spent in the nest (A), number of daily off-bouts (B) and daily mean off-bout duration in minutes (C). Colour codes represent individual nest-boxes and each point illustrates one day of incubation. Dashed black line was drawn following an intercept of 0 and a slope of 1.

These results show that the method presented here can yield accurate metrics of incubation behaviour. Based on the validation of Rhythm presented in Bueno-Enciso et al. (2017), `incR` performs better than that software and yields higher correlations between video and iButton^®^ data (Bueno-Enciso et al. 2017); however, note the possible influence of different environmental temperatures across studies. In this study the difference between nest temperatures and ambient temperatures ranged from a minimum of -0.98 (i.e. ambient temperature 0.98 degrees higher than nest temperature) to a maximum of 32.49, with a mean value across nest-boxes of 20.34°C (standard deviation = 5.18) (Table S1). For number of off-bouts, the discrepancies between `incR` and video footage seem to arise from `incR` slightly over-estimating the number of off-bouts (Figure 3B). This effect was mainly caused by data from two nest-boxes (G178_GT and GT173_GT) which were collected in the same year and location. However, the magnitude of this discrepancy was small (six off-bouts of maximum differences between estimates for whole days; estimated regression slope ± SE = 0.926 ± 0.071) and the magnitude and direction of this error is unlikely to differentially affect comparisons across groups of nests (e.g. experimental *versus* control in an experimental setup). Additional metrics to those presented here can be calculated using `incR` (Figure 1 and see package documentation), for which high reliability is expected given the results of this validation.

### Benefits of incR

The benefits of `incR` are multiple. It represents a quantitative improvement over other methods. The results of the validation suggest that `incR` may perform better than other approaches (see validation of Rhythm in Bueno-Enciso et al. 2017). No assumptions about minimum off-bout time or off-bout temperature reductions are needed and the assessment of different parameter values for `incR_scan` is straightforward (see Appendix 1 and 2). `incRscan` uses changes between consecutive temperature points, rather than total temperature reduction during an off-bout, making the detection of short off-bouts possible. Furthermore, the inclusion of data on environmental temperatures informs the analysis, allowing for off-bout detection when nest and environmental temperatures are similar. In Appendix 1 and 2, I offer detailed instructions to reproduce the analysis presented here. More generally, using a script-based approach will improve repeatability and will ease collaboration. `incR` embraces the philosophy of the R project: it is completely free and is in constant improvement. Further developments in the method to score incubation could be embedded in or used jointly with `incR` to extract metrics of incubation.

### Limitations

The capability of `incR`, or very likely of any other analytical tool to study incubation temperatures, to yield accurate results will certainly correlate with data quality. Optimal placement of the logging device among the eggs (i.e. close to the incubating adult and not buried inside nest materials) and data validation are, therefore, crucial. Two key assumptions underlie the use of `incRscan`. First, the incubating individual is assumed to rest in the nest in the `lower.time` -`upper.time` time window. This assumption holds for most species in temperate and tropical zones, for which activity outside the nest is paused during night time (a reversed pattern is expected in nocturnal species). However, careful consideration of this assumption will be needed when the species of interest do not have a rhythmic incubation pattern or rhythms differ from 24 h (Bulla et al. 2016). Secondly, the accuracy of `incRscan` will also depend on the difference between maximum incubation temperature and environmental temperature. Small differences between them will lead to subtle temperature changes after the incubating individual enters and leaves the nest, affecting the detectability of such events. The validation presented here encompasses a wide range of values for the difference between nest and environmental temperatures (Table S1) but further tests would need to be carried out to evaluate the accuracy of `incR` in hot environments. Under these conditions, apart from maximising the percentage of agreement between `incRscan` incubation scores and the data set for validation, researchers should pay careful attention to maximise agreement in other incubation metrics of such as number of incubation off-bouts. Comparing the performance of `incRscan` for data collected on the same species at different latitudes (and thus with likely large or small differences between environmental and nest temperatures) might provide valuable information on the general applicability of `incRscan`.

## Conclusions

We have developed a method that accurately extracts behavioural and temperature information from series of incubation temperature recordings. This method can potentially be used to study incubation in a broad range of species and ecological contexts and, therefore, assist the wide community of researchers studying incubation in the wild. For different species and environments, validation will be needed but we also provide detailed practical advice to carry out such validation. In order to aid its application, two appendices show in detail how researchers can easily adapt and calibrate this method to their data.

## Acknowledgements

Barbara Helm and Davide Dominoni offered the inspiration to write this manuscript: without their support and guidance none of this work would have been possible. Robyn Womack and Natalie van Dis very kindly let me use their great tit data from Glasgow and The Netherlands. They also tested the package in multiple occasions and provided thorough technical advice. Field data collection was significantly eased by the help of Davide Dominoni, Barbara Helm, Chris Pollock, Jessica Clark and Paul Jerem, along with many other volunteers. Many people at the Institute of Biodiversity, Animal Health and Comparative Medicine of the University of Glasgow provided very valuable advice; in particular, I thank Jason Matthiopoulos. I also thank Barbara Helm, Davide Dominoni, Natalie van Dis, Natalia Bravo-Santano, Andreas Nord, Caren Cooper, Wesley Hochachka and two anonymous reviewers for providing very valuable comments on an earlier draft of this manuscript.

