## Supplementary Materials for "incR: a new R package to analyse incubation behaviour"

Appendix 2 - Validating incR results

*Pablo Capilla-Lasheras*

*2018-03-21*

#### Introduction

This document provides R code to reproduce the analysis carried out in the main text of the manuscript. A brief explanation for the general coding strategy can be found in Appendix 1. The calibration of `incRscan` can be found in Appendix 1.

Change working directory and file paths according to your file system structure.

#### R code to run the incR workflow

```
# install if not done before
## for developing version
devtools::install_github(repo = "incR", username = "PabloCapilla")
## for CRAN stable version
install.packages("incR")

library(incR)
packageVersion("incR")  # release package version 1.1.0 (March 2018)

# install and load these packages to run the code below
library(doParallel)
library(ggplot2)
library(dplyr)
library(data.table)
library(extrafont)
loadfonts()
```

```
# environmenral data for Scotland
env_Data_Scotland_2015 <-
  read.csv("2-dataCalibration/envData/env_Data2015.csv")           # for 2015
env_Data_Scotland_2016 <-
  read.csv("2-dataCalibration/envData/2016/151OUT_FS_20160606.csv")# for 2016
# environmental data for the Netherlands
env_Data_NIOO_2016 <- read.csv("2-dataCalibration/envData/weathersouth_NIOO.csv")

# a sample of these data
head(env_Data_NIOO_2016)
# the other environmental temperature files have
# the same structure and column names

# nest data
incubation_files <- dir("./2-dataCalibration/data_R/", # files containing incubation data
                        full.names = TRUE,
```

```r
                         pattern = ".csv")
rawdata_incubation <- lapply(X = as.list(incubation_files), # reads every file
                                                             # in incubation_files
                               FUN = function(X) read.csv(X)) # produces a
                                                             # list of data frames

# checking the data are correct
lapply(X = rawdata_incubation, FUN = head)
```

In order from left to right: date column, nest temperature, presence of the incubating bird (1 = YES, 0 = NO) based on video footage, nest-box code, site (either Scotland or the Netherlands).

Once a list of data frames has been created, it is easy and fast to use `lapply` to apply any function to every element of the list (as done above). To make things even faster, I use `parLappy` from the `doPARALLEL` package to do parallel computation on three threads in my PC - see below how this is set up for 3 computing threads. *Doing parallel computing in R might sound difficult but it is actually very straightforward. Have a look at this* ***link*** *to find out more*

```r
ncores <- 3
cluster_incR <- makeCluster(ncores)
registerDoParallel(cluster_incR)
#getDoParWorkers() # check you actually have as many working threads as you want

clusterExport(cl = cluster_incR, c("env_Data_Scotland_2015",
                                    "env_Data_NIOO_2016",
                                    "env_Data_Scotland_2016",
                                    "rawdata_incubation"))
clusterEvalQ(cl = cluster_incR, library(incR))
clusterEvalQ(cl = cluster_incR, library(data.table))
```

Now, `parLapply` can be called. First, I use `incRprep`.

```r
# applying incRprep
incubation_prepdata <- parLapply(cl = cluster_incR,
                                  X = rawdata_incubation,
                                  fun = incRprep,
                                  date.name = "date00",
                                  temperature.name = "temp",
                                  date.format = "%d/%m/%Y %H:%M",
                                  timezone = "GMT")
```

You can check that `incRprep` has worked fine for every data frame (Note that the column *PRESENCE* gives incubation scores based on video footage). New columns `index`, `time`, `hour`, `minute`, `date`, `dec_time` and `temp1` should have been created. For information about these new variables check the documentation page for `incRprep`.

```r
lapply(incubation_prepdata, head, n = 3)
```

The next step, and final data preparation step, is to calculate environmental temperatures for every line and data frame. This task is done by `incRenv` and I apply the function in a similar fashion as we did for `incRprep`.

The only difference now is that we need to deal with the fact that the source of the environmental data changes for the different elements in `incubation_prepdata`: for the 1st and 2nd elements `env_Data_Scotland_2016` applies, `env_Data_Scotland_2015` for the 7th and 8th, and `env_Data_NIOO_2016` for 3rd to 6th one. I solve this issue in the code below, other coding choices are certainly possible.

```
clusterExport(cl = cluster_incR, c("incubation_prepdata"))
# the computation below might take between one and ten minutes
incubation_finaldata <- parLapply(cl = cluster_incR,
                                  X = as.list(c(1:length(incubation_prepdata))),
                                  fun = function(X){
                                    if(X <= 2){
                                      environmental_data <- env_Data_Scotland_2016
                                    } else {
                                      if (X >= 7){
                                        environmental_data <- env_Data_Scotland_2015
                                      } else {
                                        environmental_data <- env_Data_NIOO_2016
                                      }
                                    }
                                    output <- incRenv (data.env =
                                                         environmental_data,
                                                       data.nest =
                                                         incubation_prepdata[[X]],
                                                       env.temperature.name =
                                                         "tempEnv",
                                                       env.date.name =
                                                         "dateEnv",
                                                       env.date.format =
                                                         "%d/%m/%Y %H:%M",
                                                       env.timezone=
                                                         "GMT")
                                    return(output)
                                  }
)
# checking the new column for environmental temperatures has been added:
lapply(incubation_finaldata, head, n = 3)
```

Now we have the data ready to use `incRscan`. `incR_scan` argument values are chosen based on simulating different values and evaluating its performance (see main text for results, Appendix 1 and package vignette for detailed code to carry out such validation).

```
incRscan_list <- parLapply(cl = cluster_incR,
                           X = incubation_finaldata,
                           fun = function(X){
                             output <- incRscan(data = X,
                                                temp.name="temp",
                                                lower.time=22,
                                                upper.time=3,
                                                sensitivity = 0.25,
                                                temp.diff.threshold = 4,
                                                maxNightVariation = 1.5,
                                                env.temp = "env_temp")
                             return(output[[1]])
                           })

# checking the tables with incubation scores based on temperatures:
lapply(incRscan_list, head, 5)

# percentage of agreement between incRscan and video footage
```

```
accuracy <- parLapply(cl = cluster_incR,
                      X = incRscan_list,
                      fun = function (X) {
                        if(is.null(dim(X))){
                          return(NA)
                        } else {
                          X <- X[complete.cases(X$PRESENCE),]
                          1 - (sum(abs(X$incR_score-X$PRESENCE)) / length(X$incR_score))
                        }
                      }

)

# summary of accuracy
summary(do.call(what = "rbind", args = accuracy))
```

#### Correlations between video-based and incR-based incubation attendance

```
attendance_results <- parLapply(cl = cluster_incR,
                                X = incRscan_list,
                                fun = function(X){
                                  X <- X[complete.cases(X$PRESENCE),]
                                  output_video <-
                                    incRatt(data = X,
                                            vector.incubation = "PRESENCE")
                                  output_video$box <-
                                    rep(na.omit(unique(X$BOX)),
                                        length = nrow(output_video))
                                  output_incR <-
                                    incRatt(data = X,
                                            vector.incubation = "incR_score")
                                  output_video$incR_scan <- output_incR[,2]
                                  return(output_video)
                                })

attendance_results <- rbindlist(attendance_results)
cor.test(attendance_results$percentage_in, attendance_results$incR_scan)

attendance_plot <- ggplot(data = attendance_results,
                          aes(x = incR_scan, y = percentage_in, color = box)) +
  geom_point(size = 7) +
  geom_abline(slope = 1, intercept = 0, linetype = 2, size = 1.5) +
  theme_bw() +
  labs(x = " ", y = " ") +
  theme(axis.title.x = element_text(family = "Courier New",
                                    colour="black", size=35),
        axis.text.x  = element_text(size=25),
        axis.title.y = element_text(family = "Times New Roman", colour="black", size=35,
                                    margin = margin (0,25,0,0)),
        axis.text.y  = element_text(size=25),
```

```
        panel.grid.minor.x=element_blank(),
        panel.grid.minor.y=element_blank(),
        panel.grid.major.y=element_blank(),
        panel.grid.major.x=element_blank(),
        legend.position="none")
```

#### Correlations between video-based and incR-based number and duration of off-bouts

```
offbout_results <- parLapply(cl = cluster_incR,
                             X = incRscan_list,
                             fun = function(X){
                               X <- X[complete.cases(X$PRESENCE),]
                               sampling_rate <- X$dec_time[5] - X$dec_time[4]

                               output_bouts <- {
                                 incRbouts(data = X,
                                           vector.incubation = "PRESENCE",
                                           dec_time = "dec_time",
                                           temp = "temp",
                                           sampling.rate = sampling_rate)
                               }$day_bouts
                               output_bouts$box <- rep(na.omit(unique(X$BOX)),
                                                       length = nrow(output_bouts))

                               output_bouts$incR_scan_nbouts <- {
                                 incRbouts(data = X,
                                           vector.incubation = "incR_score",
                                           dec_time = "dec_time",
                                           temp = "temp",
                                           sampling.rate = sampling_rate)}$day_bouts[,3]
                               output_bouts$incR_scan_timebouts <- {
                                 incRbouts(data = X,
                                           vector.incubation = "incR_score",
                                           dec_time = "dec_time",
                                           temp = "temp",
                                           sampling.rate = sampling_rate)}$day_bouts[,5]

                               return(output_bouts)
                             })



bouts_results <- rbindlist(offbout_results)
cor.test(bouts_results$number.off.bouts,bouts_results$incR_scan_nbouts)
cor.test(bouts_results$mean.time.off.bout,bouts_results$incR_scan_timebouts)


nbout_plot <- ggplot(data = bouts_results,
                     aes(x = incR_scan_nbouts, y = number.off.bouts, color = box)) +
  geom_point(size = 7) +
```

```r
  geom_abline(slope = 1, intercept = 0, linetype = 2, size = 1.5) +
  theme_bw() +
  labs(x = " ", y = "Video-footage estimate") +
  theme(axis.title.x = element_text(family = "Courier New",
                                    colour="black", size=35),
        axis.text.x  = element_text(size=25),
        axis.title.y = element_text(family = "Times New Roman", colour="black", size=35,
                                    margin = margin (0,25,0,0)),
        axis.text.y  = element_text(size=25),
        panel.grid.minor.x=element_blank(),
        panel.grid.minor.y=element_blank(),
        panel.grid.major.y=element_blank(),
        panel.grid.major.x=element_blank(),
        legend.position="none")

time_bout_plot <- ggplot(data = bouts_results,
       aes(x = incR_scan_timebouts*60, y = mean.time.off.bout*60, color = box)) +
  geom_point(size = 7) +
  geom_abline(slope = 1, intercept = 0, linetype = 2, size = 1.5) +
  theme_bw() +
  labs(x = "incR estimate", y = " ") +
  theme(axis.title.x = element_text(family = "Courier New",
                                    colour="black", size=35),
        axis.text.x  = element_text(size=25),
        axis.title.y = element_text(family = "Times New Roman", colour="black", size=35,
                                    margin = margin (0,25,0,0)),
        axis.text.y  = element_text(size=25),
        panel.grid.minor.x=element_blank(),
        panel.grid.minor.y=element_blank(),
        panel.grid.major.y=element_blank(),
        panel.grid.major.x=element_blank(),
        legend.position="bottom",
        legend.title = element_blank(),
        legend.text = element_text(family = "Times New Roman",
                                   size = 20),
        legend.key = element_rect(size = 5),
        legend.key.size = unit(1.7, 'lines'))

# creating the plot panel
source("http://peterhaschke.com/Code/multiplot.R")
ggsave(plot = multiplot(plotlist = list(attendance_plot, nbout_plot, time_bout_plot),
         cols = 1), filename = "./plots/Figure 3.jpeg",
       device = "jpeg",
       height = 15, width = 10)
```

#### Creating nest temperature traces

As an visual example of what the package performs, I represent the nest temperature trace of a nest box at different stages of the analysis.

```r
inc_trace1 <- ggplot(data = incRscan_list[[8]] %>%
                       filter(date == "2015-05-07"),
```

```
                    aes(x = dec_time, y = temp)) +
  geom_point(size = 1.75, color = "grey") +
  theme_bw() +
  labs(y = "Temperature (ºC)",
       x = "Time") +
  scale_color_manual(name = "", labels = c("Absence", "Presence"),
                    values = c("#8da0cb", "#fc8d62")) +
  theme(axis.title.x = element_text(family = "Times New Roman",
                                    colour="black", size=20),
        axis.text.x  = element_text(size=17, family = "Times New Roman"),
        axis.title.y = element_text(family = "Times New Roman",
                                    colour="black", size=18,
                                    margin = margin (0,5,0,0)),
        axis.text.y  = element_text(size=17,family = "Times New Roman"),
        panel.grid.minor.x=element_blank(),
        panel.grid.minor.y=element_blank(),
        panel.grid.major.y=element_blank(),
        panel.grid.major.x=element_blank()) +
  scale_y_continuous(breaks = seq(10,38,4),
                     labels = seq(10,38,4)) +
  scale_x_continuous(breaks = seq(2,24,2),
                     labels = seq(2,24,2))

ggsave(filename = "./plots/inc_trace1.jpeg", plot = inc_trace1,
       device = "jpeg", width = 5, height = 5)

inc_trace2 <- ggplot(data = incRscan_list[[8]] %>%
                      filter(date == "2015-05-07"),
                    aes(x = dec_time, y = temp)) +
  geom_point(size = 1.75, color = "grey") +
  geom_line(aes(y = env_temp), color = "#66c2a5", size = 1.5) +
  theme_bw() +
  labs(y = "Temperature (ºC)",
       x = "Time") +
  scale_color_manual(name = "", labels = c("Absence", "Presence"),
                    values = c("#8da0cb", "#fc8d62")) +
  theme(axis.title.x = element_text(family = "Times New Roman",
                                    colour="black", size=20),
        axis.text.x  = element_text(size=17, family = "Times New Roman"),
        axis.title.y = element_text(family = "Times New Roman",
                                    colour="black", size=18,
                                    margin = margin (0,5,0,0)),
        axis.text.y  = element_text(size=17,family = "Times New Roman"),
        panel.grid.minor.x=element_blank(),
        panel.grid.minor.y=element_blank(),
        panel.grid.major.y=element_blank(),
        panel.grid.major.x=element_blank()) +
  scale_y_continuous(breaks = seq(10,38,4),
                     labels = seq(10,38,4)) +
  scale_x_continuous(breaks = seq(2,24,2),
                     labels = seq(2,24,2))

ggsave(filename = "./plots/inc_trace2.jpeg", plot = inc_trace2,
```

```
       device = "jpeg", width = 5, height = 5)


inc_trace3 <- ggplot(data = incRscan_list[[8]] %>%
                       filter(date == "2015-05-07"),
                     aes(x = dec_time, y = temp, color = factor(incR_score)))+
  geom_point(size = 1.75) +
  geom_line(aes(y = env_temp), color = "#66c2a5", size = 1.5) +
  theme_bw() +
  labs(y = "Temperature (ºC)",
       x = "Time") +
  scale_color_manual(name = "", labels = c("Absence", "Presence"),
                     values = c("#8da0cb", "#fc8d62")) +
  theme(axis.title.x = element_text(family = "Times New Roman",
                                     colour="black", size=20),
        axis.text.x  = element_text(size=17, family = "Times New Roman"),
        axis.title.y = element_text(family = "Times New Roman",
                                     colour="black", size=18,
                                     margin = margin (0,5,0,0)),
        axis.text.y  = element_text(size=17,family = "Times New Roman"),
        panel.grid.minor.x=element_blank(),
        panel.grid.minor.y=element_blank(),
        panel.grid.major.y=element_blank(),
        panel.grid.major.x=element_blank(),
        legend.position="top",
        legend.text = element_text(family = "Times New Roman", size = 17))+
  scale_y_continuous(breaks = seq(10,38,4),
                      labels = seq(10,38,4)) +
  scale_x_continuous(breaks = seq(2,24,2),
                      labels = seq(2,24,2))

ggsave(filename = "./plots/inc_trace3.jpeg", plot = inc_trace3,
       device = "jpeg", width = 5, height = 5)
```

### Supplementary Figure 2

```
SupFig2 <- incRplot(data = data %>%
                      filter(BOX == "G16_BT") %>%
                      filter(date == "2015-05-09"),
                    time.var = "dec_time",
                    day.var = "date",
                    inc.temperature.var = "temp",
                    env.temperature.var = "env_temp",
                    vector.incubation = "incR_score")

SupFig2 <- SupFig2 +
  theme(axis.title.x = element_text(family = "Times New Roman",
                                     colour="black", size=20),
        axis.text.x  = element_text(size=17, family = "Times New Roman"),
        axis.title.y = element_text(family = "Times New Roman",
                                     colour="black", size=18,
```

```
                                    margin = margin (0,5,0,0)),
        axis.text.y  = element_text(size=17,family = "Times New Roman"),
        panel.grid.minor.x=element_blank(),
        panel.grid.minor.y=element_blank(),
        panel.grid.major.y=element_blank(),
        panel.grid.major.x=element_blank(),
        legend.position="top",
        legend.text = element_text(family = "Times New Roman", size = 17))

ggsave(filename = "./plots/SupFig2.jpeg", plot = SupFig2,
       device = "jpeg", width = 7, height = 5)
```
