## Supplementary Materials for "incR: a new R package to analyse incubation behaviour"

**Supplementary Figures**

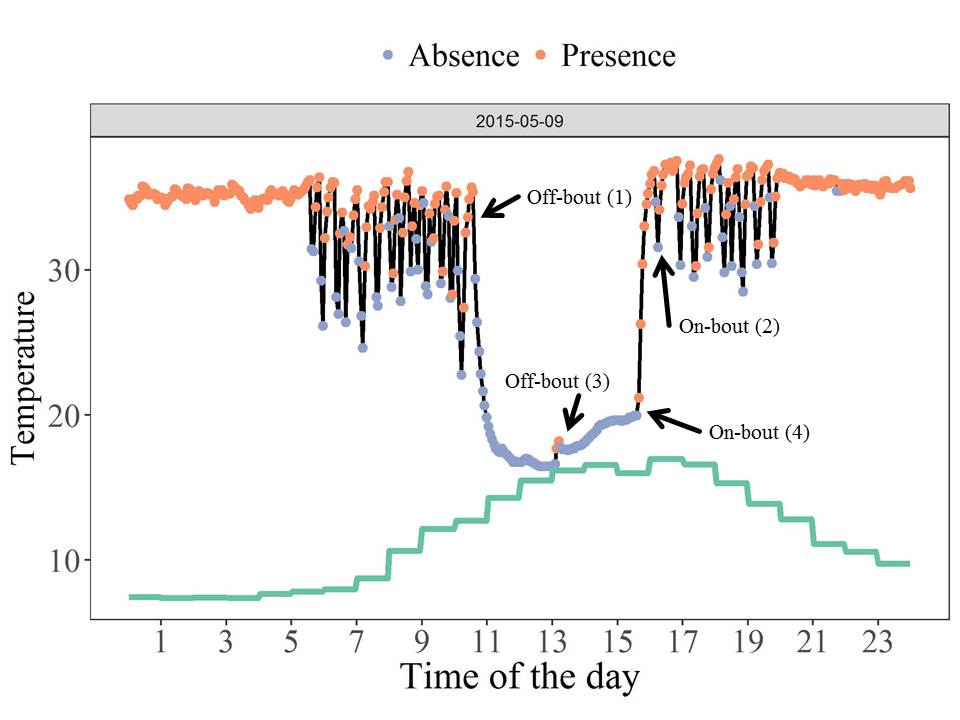

**Figure S1.** One day of incubation temperatures (dots connected by black line) for nest-box G16_BT. incR provides the function incRplot to generate this plot, representing on-bouts (light-red) and off-bouts (purple) along with environmental temperatures (light-green line). Two on-bouts and off-bouts have been marked with arrows to represent examples of scenarios 1, 2, 3 and 4 (numbers in brackets) as explained in the “*Automated incubation scoring:* incRscan” section of the main text.

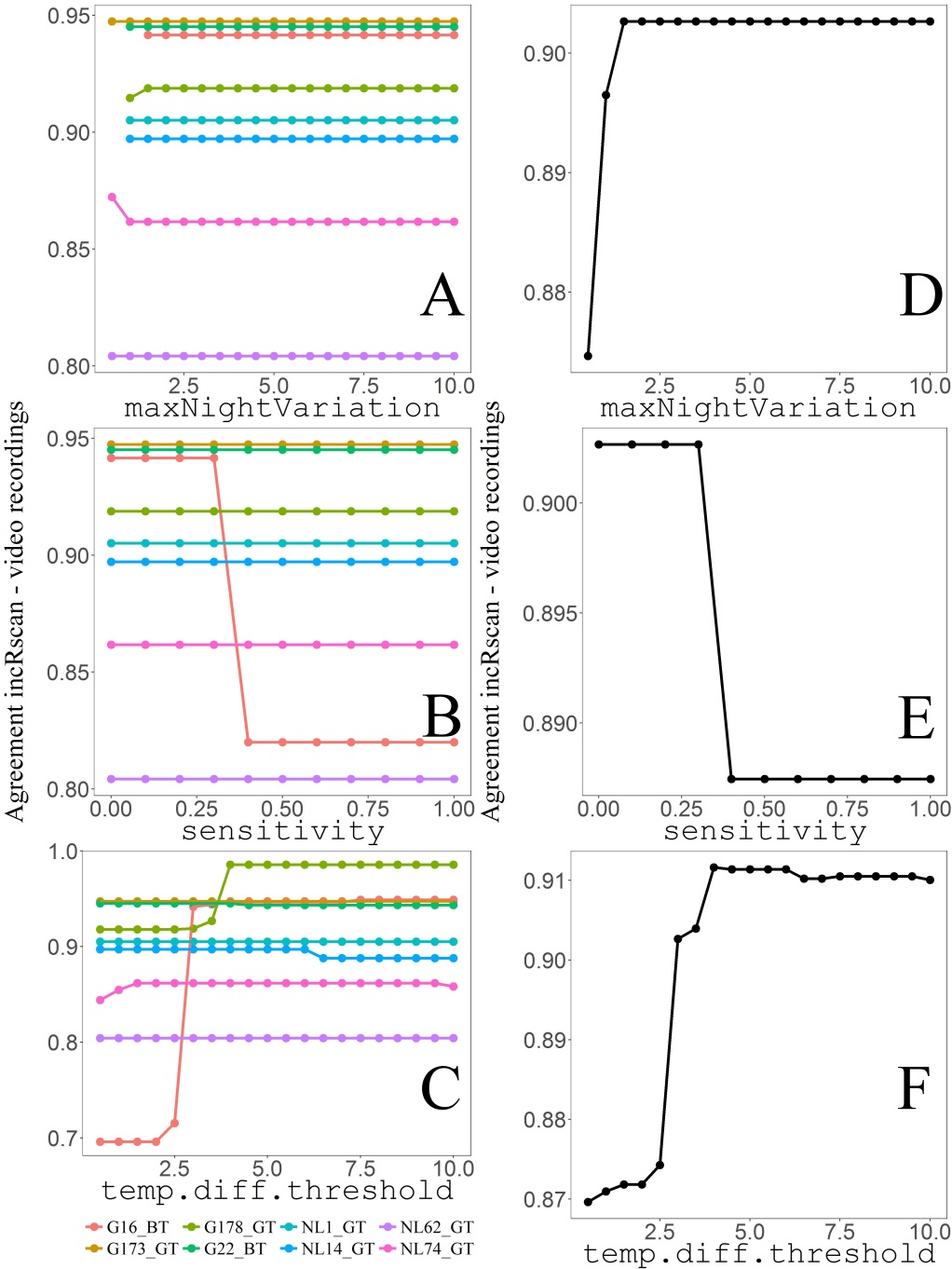

**Figure S2**. Results of the 1-dimensinal grid search for individual nest-boxes (A, B and C) and averaging over nest-boxes (D, E, F). Plots show the percentage of agreement between incRscan-based and video-based incubation scores after varying values of maxNightVariation (A, D), sensitivity (B, E), temp.diff.threshold (C, F).

**Supplementary Tables**

**Table S1.** Minimum, mean and maximum values (in °C) for the difference between nest and environmental temperatures across nest-boxes.

| **Nest-box** | **Minimum** | **Mean** | **Maximum** |
| --- | --- | --- | --- |
| G173_GT | 17.49 | 27.22 | 32.49 |
| G178_GT | 9.71 | 23.18 | 31.83 |
| NL1_GT | 8.44 | 17.88 | 19.92 |
| NL14_GT | 0.73 | 14.56 | 18.31 |
| NL62_GT | 10.61 | 16.82 | 19.44 |
| NL74_GT | -0.98 | 13.80 | 19.41 |
| G16_BT | 1.50 | 25.33 | 28.50 |
| G22_BT | 4.04 | 23.94 | 27.39 |
