## Supplementary Materials for "incR: a new R package to analyse incubation behaviour"

Appendix 1

*Pablo Capilla-Lasheras*

### Introduction

The use of the main function in `incR`, `incRscan`, requires to set argument values that can affect `incRscan` performance. The combination of video-recordings and nest temperatures can provide a powerful way of (i) assessing the general performance of `incRscan` on a specific data set and (ii) simulating how robust the results are to varying `incRscan` parameters, enabling the user to choose the optimal settings.

Here, using incubation temperatures from blue tits (*Cyanistes caeruleus*) and great tits (*Parus major*) recorded by iButtons (Maxim Technologies Inc.), I apply the approach described above. I combine nest temperatures and video-recordings to evaluate the position of the incubating individual (in or out the nest - incubation scores), and find the optimal parameters for `incRscan` for each data set.

#### The data

See the main manuscript for details about data collection. I analyse twelve (total or partial) days of incubation coming from eight different nest-boxes (two blue tits and six great tit clutches) in two years of study, for which video-cameras were simultaneously employed along with iButtons devices that recorded nest temperature. The data were collected in spring 2015 and 2016 in Scotland and The Netherlands.

### Workflow

Following the recommendations in the main manuscript - that can also be found in an `incR` vignette:

### Data preparation `incRprep` and `incRenv`

I load data from ten nest-boxes as ten different files and then create a list in which every element is a data frame. Of course, if you wanted to reproduce the code using your own data, you will need to change the paths to files.

```
# install if not done before
## for developing version
devtools::install_github(repo = "PabloCapilla/incR")
## for CRAN stable version
install.packages("incR")

library(incR)
packageVersion("incR")  # release package version 1.1.0 (March 2018)


# install and load this packages to run this vignette
library(ggplot2)
library(dplyr)
library(data.table)
```

```r
library(extrafont)
loadfonts()
```

```r
# environmenral data for Scotland for 2015 and  2016
env_Data_Scotland_2015 <- read.csv("2-dataCalibration/envData/env_Data2015.csv")
env_Data_Scotland_2016 <- read.csv("2-dataCalibration/envData/2016/1510UT_FS_20160606.csv")
# environmental data for the Netherlands
env_Data_NIOO_2016 <- read.csv("2-dataCalibration/envData/weathersouth_NIOO.csv")

# a sample of these data
head(env_Data_NIOO_2016) # the other env files have the same structure and column names

# incubation data
incubation_files <- dir("./2-dataCalibration/data_R/", # files containing incubation data
                        full.names = TRUE,
                        pattern = ".csv")
rawdata_calibration <- lapply(X = as.list(incubation_files), # reads every file
                                                             # in incubation_files
                              FUN = function(X) read.csv(X)) # produces a list
                                                             # of data frames

# checking the data are correct
lapply(X = rawdata_calibration, FUN = head)
```

In order from left to right: date column, nest temperature, presence of the incubating bird (1 = YES, 0 = NO, or NA) based on video footage, nest-box code, site (either Scotland or the Netherlands).

Once a list of data frames has been created, it is easy and fast to use `lapply` to apply any function to every element of the list (as done above). To make things even faster, I use `parLappy` from the `doPARALLEL` package to do parallel computation on three threads in my PC - see below how this is set up for 3 computing threads. *Doing parallel computing in R might sound difficult but it is actually very straightforward. Have a look at this* ***link*** *to find out more*

```r
library(doParallel)
ncores <- 3
cluster_incR <- makeCluster(ncores)
registerDoParallel(cluster_incR)
#getDoParWorkers() # check you actually have as many working threads as you want

clusterExport(cl = cluster_incR, c("env_Data_Scotland_2015",
                                   "env_Data_NIOO_2016",
                                   "env_Data_Scotland_2016",
                                   "rawdata_calibration"))
clusterEvalQ(cl = cluster_incR, library(incR))
```

Now, `parLapply` can be called.

```r
# applying incRprep
calibration_prepdata <- parLapply(cl = cluster_incR,
                                  X = rawdata_calibration,
                                  fun = incRprep,
                                  date.name = "date00",
                                  temperature.name = "temp",
                                  date.format = "%d/%m/%Y %H:%M",
                                  timezone = "GMT")
```

You can check that `incRprep` has worked fine for every data frame (Note that the column *PRESENCE* gives incubation scores based on video footage). New columns `index`, `time`, `hour`, `minute`, `date`, `dec_time` and `temp1` should have been created. For information about these new variables check the documentation page for `incRprep`.

```
lapply(calibration_prepdata, head, n = 3)
```

The next step, and final data preparation step, is to calculate environmental temperatures for every line and data frame. This task is done by `incRenv` and I apply the function in a similar fashion as we did for `incRprep`.

The only difference now is that we need to deal with the fact that the source of the environmental data changes for the different elements in `calibration_prepdata`: for the 1st and 2nd elements `env_Data_Scotland_2016` applies, `env_Data_Scotland_2015` for the 7th and 8th, and `env_Data_NIOO_2016` for 3rd to 6th one. I solve this issue in the code below, other coding choices are certainly possible.

```
clusterExport(cl = cluster_incR, c("calibration_prepdata"))
```

```
# the computation below might take between one and ten minutes
calibration_finaldata <- parLapply(cl = cluster_incR,
                                   X = as.list(c(1:length(calibration_prepdata))),
                                   fun = function(X){
                                     if(X <= 2){
                                       environmental_data <- env_Data_Scotland_2016
                                     } else {
                                       if (X >= 7){
                                         environmental_data <- env_Data_Scotland_2015
                                       } else {
                                         environmental_data <- env_Data_NIOO_2016
                                       }
                                     }
                                     output <- incRenv (data.env = environmental_data,
                                                        data.nest =
                                                          calibration_prepdata[[X]],
                                                        env.temperature.name = "tempEnv",
                                                        env.date.name = "dateEnv",
                                                        env.date.format =
                                                          "%d/%m/%Y %H:%M",
                                                        env.timezone="GMT")
                                     return(output)
                                   }
)
#stopCluster(cluster_incR)   # stop cluster
# checking the new column for environmental temperatures has been added:
lapply(calibration_finaldata, head, n = 3)

# range in differences between nest temperature and environmental temperature
temp_dif <- rbindlist(lapply(calibration_finaldata, function(x){
  df <- x %>% filter(PRESENCE == 1)                    # select only data used for calibration
  data.frame(box = unique(x$BOX),
             mean = mean(df$temp - df$env_temp),
             min = range(df$temp - df$env_temp)[1],     # calculate minimum for every nest-box
             max = range(df$temp - df$env_temp)[2])     # calculate maximum for every nest-box
})
)
```

```r
# mean across nest-boxes
mean(temp_dif$mean)
# sd
sd(temp_dif$mean)
```

Now we have the data ready to use `incRscan` and assess its results over a number of different parameter values.

### Calibration of `incRscan`

For details about the parameters in `incRscan`, check the documention (`?incR::incRscan`) and the main manuscript.

I run `incRscan` for varying values of `maxNightVariation`, `sensitivity` and `temp.diff.threshold`, as specified in the main text, while keeping the other parameters to fixed values: `lower.time` = 22pm, `upper.time` = 3am, `maxNightVariation` = 1.5, `sensitivity` = 0.15, `temp.diff.threshold` = 3.

I simply set parameter values for testing, by building a matrix that contains every single combination of parameter values under inspection and run `incRscan` for each of those rows (i.e. for every parameter combination). Then, I compare `incRscan`-based incubation scores (stored in a new column `incR_score`) with video-based incubation scores (in `PRESENCE`).

#### Setting up matrix of parameter values:

```r
# parameter values for simulation
sim.maxNightVariation <- seq (from = 0.5, to = 10, by = 0.5)
sim.sensitivity <- seq (from = 0, to = 1, by = 0.1)
sim.temp.diff.threshold <- seq (from = 0.5, to = 10, by = 0.5)

# parameter combinations for maxNightVariation
sim.maxNightVariation_table <- data.frame(sim_maxNightVariation = sim.maxNightVariation,
                                          sim_sensitivity = 0.15,
                                          sim_temp.diff.threshold = 3,
                                          batch = 1)
# parameter combinations for sensitivity
sim.sensitivity_table <- data.frame(sim_maxNightVariation = 1.5,
                                    sim_sensitivity = sim.sensitivity,
                                    sim_temp.diff.threshold = 3,
                                    batch = 2)
# parameter combinations for temp.diff.threshold
sim.temp.diff.threshold_table <- data.frame(sim_maxNightVariation = 1.5,
                                  sim_sensitivity = 0.15,
                                  sim_temp.diff.threshold = sim.temp.diff.threshold,
                                  batch = 3)
# table for every parameter combination of interest
parameter_table <- rbind(sim.maxNightVariation_table,
                         sim.sensitivity_table,
                         sim.temp.diff.threshold_table)

head(parameter_table)
nrow(parameter_table) # meaning that incRscan need to run 52 times
```

```
clusterExport(cl = cluster_incR, c("parameter_table", "calibration_finaldata"))
```

##### Running `incRscan` for every combination of parameter values

As well as running `incRscan`, I compute the percentage of agreement between `incR_score` and `PRESENCE` for every iteration (i.e. parameter combination).

```
progressBar <- txtProgressBar(min = 0, max = nrow(parameter_table), style = 3) # use in PC
accuracy_results <- as.list(NA) # list to store results

for (i in 1:nrow(parameter_table)){
  clusterExport(cl = cluster_incR, c("i"))
  incRscan_list <- parLapply(cl = cluster_incR,
                             X = calibration_finaldata,
                             fun = function(X){
                               output <- incRscan(
                                 data = X,
                                 temp.name="temp",
                                 lower.time=22,
                                 upper.time=3,
                                 sensitivity =
                                   parameter_table[i,"sim_sensitivity"],
                                 temp.diff.threshold =
                                   parameter_table[i,"sim_temp.diff.threshold"],
                                 maxNightVariation =
                                   parameter_table[i,"sim_maxNightVariation"],
                                 env.temp = "env_temp"
                               )
                               return(output[[1]])
                             })
  # percentage of agreement between incRscan and video footage
  accuracy <- parLapply(cl = cluster_incR,
                        X = incRscan_list,
                        fun = function (X) {
                          if(is.null(dim(X))){
                            return(NA)
                          } else {
                            X <- X[complete.cases(X$PRESENCE),]
                            1 - (sum(
                              abs(X$incR_score-X$PRESENCE)) / length(X$incR_score)
                            )
                          }
                        }

  )
  # results
  accuracy_results[[i]] <- data.frame(box = do.call(
    args = lapply(X = calibration_finaldata,
                  FUN = function(X) as.character(unique(X$BOX))),
    what = "rbind"),
    sensitivity = rep(parameter_table[i,"sim_sensitivity"],8),
    temp.diff.threshold = rep(parameter_table[i,"sim_temp.diff.threshold"],8),
    maxNightVariation = rep(parameter_table[i,"sim_maxNightVariation"],8),
```

```
    batch = rep(parameter_table[i,"batch"],8),
    accuracy = do.call(args = accuracy,
                       what = "rbind")
  )

  setTxtProgressBar(pb = progressBar, value = i)
}

# bind list in a data frame
calibrating_results <- rbindlist(accuracy_results)
```

### Visualising results of the calibration

Once the loop finishes the calculation, we can visualise which combination of parameters provides the highest agreement between incubation scores based on video footage and `incRscan`.

For simulation of `maxNightVariation` values:

```
calibration_plot1 <- ggplot(data = calibrating_results %>% filter(batch == 1),
                            aes(x = maxNightVariation, y = accuracy, color = box))+
  geom_point(size = 4) +
  geom_line(size = 1.5) +
  labs(x = "maxNightVariation", y = " ")+
  theme_bw() +
  theme(axis.title.x = element_text(family = "Courier New",
                                    colour="black", size=35),
        axis.text.x  = element_text(size=25),
        axis.title.y = element_text(family = "Times New Roman", colour="black", size=35,
                                    margin = margin (0,25,0,0)),
        axis.text.y  = element_text(size=25),
        panel.grid.minor.x=element_blank(),
        panel.grid.minor.y=element_blank(),
        panel.grid.major.y=element_blank(),
        panel.grid.major.x=element_blank(),
        legend.position="none",
        legend.text = element_text(family = "Times New Roman",
                                   size = 17))
```

For simulation of `sensitivity` values:

```
calibration_plot2 <- ggplot(data = calibrating_results %>% filter(batch == 2),
                            aes(x = sensitivity, y = accuracy, color = box))+
  geom_point(size = 4) +
  geom_line(size = 1.5) +
  theme_bw() +
  labs(x = "sensitivity", y = "Agreement incRscan - video recordings")+
  theme(axis.title.x = element_text(family = "Courier New",
                                    colour="black", size=35),
        axis.text.x  = element_text(size=25),
        axis.title.y = element_text(family = "Times New Roman", colour="black", size=30,
                                    margin = margin (0,25,0,0)),
        axis.text.y  = element_text(size=25),
        panel.grid.minor.x=element_blank(),
```

```
        panel.grid.minor.y=element_blank(),
        panel.grid.major.y=element_blank(),
        panel.grid.major.x=element_blank(),
        legend.position="none",
        legend.text = element_text(family = "Times New Roman",
                                   size = 17))
```

For simulation of `temp.diff.threshold` values:

```
calibration_plot3 <- ggplot(data = calibrating_results %>% filter(batch == 3),
       aes(x = temp.diff.threshold, y = accuracy, color = box))+
  geom_point(size = 4) +
  geom_line(size = 1.5) +
  theme_bw() +
  labs(x = "temp.diff.threshold", y = " ")+
  theme(axis.title.x = element_text(family = "Courier New",
                                    colour="black", size=35),
        axis.text.x  = element_text(size=25),
        axis.title.y = element_text(family = "Times New Roman", colour="black", size=35,
                                    margin = margin (0,25,0,0)),
        axis.text.y  = element_text(size=25),
        panel.grid.minor.x=element_blank(),
        panel.grid.minor.y=element_blank(),
        panel.grid.major.y=element_blank(),
        panel.grid.major.x=element_blank(),
        legend.position="bottom",
        legend.title = element_blank(),
        legend.text = element_text(family = "Times New Roman",
                                   size = 20),
        legend.key = element_rect(size = 5),
        legend.key.size = unit(1.7, 'lines'))
```

Averaging over nest-boxes for each set of simulations, `maxNightVariation` values above 1.5, `sensitivity` between 0 and 0.3 and `temp.diff.threshold` of 4 yielded the highest percentage of agreement between `incRscan` and video recordings (90.27%, 90.27% and 91.16% respectiuvely (Figure below).

```
# for maxNightVariation
calibration_plot1_mean <- calibrating_results %>%
  filter(batch == 1) %>%
  group_by(maxNightVariation) %>%
  summarise(mean_accuracy = mean(accuracy, na.rm = TRUE)) %>%
  arrange(desc(mean_accuracy)) %>%
  ggplot(aes(y = mean_accuracy, x = maxNightVariation)) +
  geom_point(size = 4) +
  geom_line(size = 1.5)+
  theme_bw() +
  labs(x = "maxNightVariation", y = " ")+
  theme(axis.title.x = element_text(family = "Courier New",
                                    colour="black", size=35),
        axis.text.x  = element_text(size=25),
        axis.title.y = element_text(family = "Times New Roman", colour="black", size=35,
                                    margin = margin (0,25,0,0)),
        axis.text.y  = element_text(size=25),
        panel.grid.minor.x=element_blank(),
        panel.grid.minor.y=element_blank(),
```

```
        panel.grid.major.y=element_blank(),
        panel.grid.major.x=element_blank(),
        legend.position="none",
        legend.title = element_blank(),
        legend.text = element_text(family = "Times New Roman",
                                   size = 17))
# for sensitivity
calibration_plot2_mean <- calibrating_results %>%
  filter(batch == 2) %>%
  group_by(sensitivity) %>%
  summarise(mean_accuracy = mean(accuracy, na.rm = TRUE)) %>%
  arrange(desc(mean_accuracy)) %>%
  ggplot(aes(y = mean_accuracy, x = sensitivity)) +
  geom_point(size = 4) +
  geom_line(size = 1.5)+
  theme_bw() +
  labs(x = "sensitivity", y = "Agreement incRscan - video recordings")+
  theme(axis.title.x = element_text(family = "Courier New",
                                    colour="black", size=35),
        axis.text.x  = element_text(size=25),
        axis.title.y = element_text(family = "Times New Roman", colour="black", size=30,
                                    margin = margin (0,25,0,0)),
        axis.text.y  = element_text(size=25),
        panel.grid.minor.x=element_blank(),
        panel.grid.minor.y=element_blank(),
        panel.grid.major.y=element_blank(),
        panel.grid.major.x=element_blank(),
        legend.position="none",
        legend.title = element_blank(),
        legend.text = element_text(family = "Times New Roman",
                                   size = 30))
# for temp.diff.threshold
calibration_plot3_mean <- calibrating_results %>%
  filter(batch == 3) %>%
  group_by(temp.diff.threshold) %>%
  summarise(mean_accuracy = mean(accuracy, na.rm = TRUE)) %>%
  arrange(desc(mean_accuracy)) %>%
  ggplot(aes(y = mean_accuracy, x = temp.diff.threshold)) +
  geom_point(size = 4)+
  geom_line(size = 1.5)+
  theme_bw() +
  labs(x = "temp.diff.threshold", y = " ")+
  theme(axis.title.x = element_text(family = "Courier New",
                                    colour="black", size=35),
        axis.text.x  = element_text(size=25),
        axis.title.y = element_text(family = "Times New Roman", colour="black", size=35,
                                    margin = margin (0,25,0,0)),
        axis.text.y  = element_text(size=25),
        panel.grid.minor.x=element_blank(),
        panel.grid.minor.y=element_blank(),
        panel.grid.major.y=element_blank(),
        panel.grid.major.x=element_blank(),
        legend.position="none",
```

```
        legend.title = element_blank(),
        legend.text = element_text(family = "Times New Roman",
                                   size = 17))

# plot to visualise best results per test per box
best_agreement <- calibrating_results %>%
  group_by(box, batch) %>%
  arrange(desc(accuracy)) %>%
  slice(1) %>%
  ggplot(aes(x = box, y = accuracy, fill = box)) +
  labs(x = "Nest-box", y = "Agreement incRscan - video recordings")+
  geom_point(position = position_jitter(width = 0.4),
             color = "black",
             pch = 21,
             alpha = 0.6,
             size = 6.5) +
  theme_bw()+
  theme(axis.title.x = element_text(family = "Times New Roman",
                                    colour="black",
                                    size=35),
        axis.text.x  = element_text(size=25,
                                    angle = 45,
                                    family = "Times New Roman"),
        axis.title.y = element_text(family = "Times New Roman",
                                    colour="black",
                                    size=35,
                                    margin = margin (0,25,0,0)),
        axis.text.y  = element_text(size=25),
        panel.grid.minor.x=element_blank(),
        panel.grid.minor.y=element_blank(),
        panel.grid.major.y=element_blank(),
        panel.grid.major.x=element_blank(),
        legend.position="none",
        legend.title = element_blank(),
        legend.text = element_text(family = "Times New Roman",
                                   size = 35))


ggsave(plot = best_agreement,
       file = "./plots/Figure 2.jpeg",
       device = "jpeg",
       height = 11, width = 13)

# creating the panel
# plotting the six plots together
source("http://peterhaschke.com/Code/multiplot.R")
ggsave(plot = multiplot(plotlist = list(calibration_plot1,
                                        calibration_plot2,
                                        calibration_plot3,
                                        calibration_plot1_mean,
                                        calibration_plot2_mean,
                                        calibration_plot3_mean),
                        cols = 2),
```

```
       filename = "./plots/Supplementary_Figure_1.jpeg",
       device = "jpeg",
       height = 20, width = 15)
```
